# Evaluation of natural killer cell tumor homing and effector function in response to CDK4/6 and AURKA inhibition in a melanoma tumor-on-a-chip platform

**DOI:** 10.1101/2025.07.09.662476

**Authors:** Srija Chakraborty, Connor Durham, Vijaya Bharti, Marina Capece, Alexander E. Davies, Anna Vilgelm, Aleksander Skardal

**Author notes:** Correspondence: Aleksander Skardal, PhD, 2107 Pelotonia Reseach Center, 2255 Kenny Road, Columbus, Ohio 43210.

## Abstract

Natural killer (NK) cells have emerged as an important clinical tool cellular immunotherapy. Whereas immune checkpoint blockade (ICB) or chimeric antigen receptor (CAR) T-cell therapy (CAR-T) therapy have been adopted as a first line treatments in different malignancies, such as melanoma, these approaches do not work for all patients. T cells require proper antigen presentation on tumor cells for recognition and to carry out their corresponding cytotoxic functions. Deficiency of tumor antigens, or high variability in those present, make T cell-based CAR-T and ICB ineffective. By contrast, NK cells are not limited by antigen presentation deficiencies, offering a potential alternative approach, yet their efficacy can suffer from immunosuppressive signals. Herein, we sought to develop in vitro and on-chip platforms to identify strategies for enhance, rather than suppress, NK cell homing to tumor cells. We explored the use of inhibition of kinases such as CK4/6 and AURKA to induce tumor cell production of chemokines that NK cells migrate towards in aggressive melanoma models. We evaluated chemokine-aided NK cell migration-homing capabilities and their therapeutic efficacy and found that treatment of both melanoma cell line and patient-tumor constructs (PTCs) with CDK4/6 and AURKA generally resulted in improved NK cell homing to tumor cells and accompanying tumor cell killing. Interestingly, this chemokine-guided NK cell migration did not generate as effective outcomes in models using a mildly aggressive melanoma cell line. For our studies, we used 3D tumor constructs in both static Transwell models and then in a bioengineered NK cell-functionalized tumor-on-a-chip (NK-TOC) platform.

**Graphical abstract:** 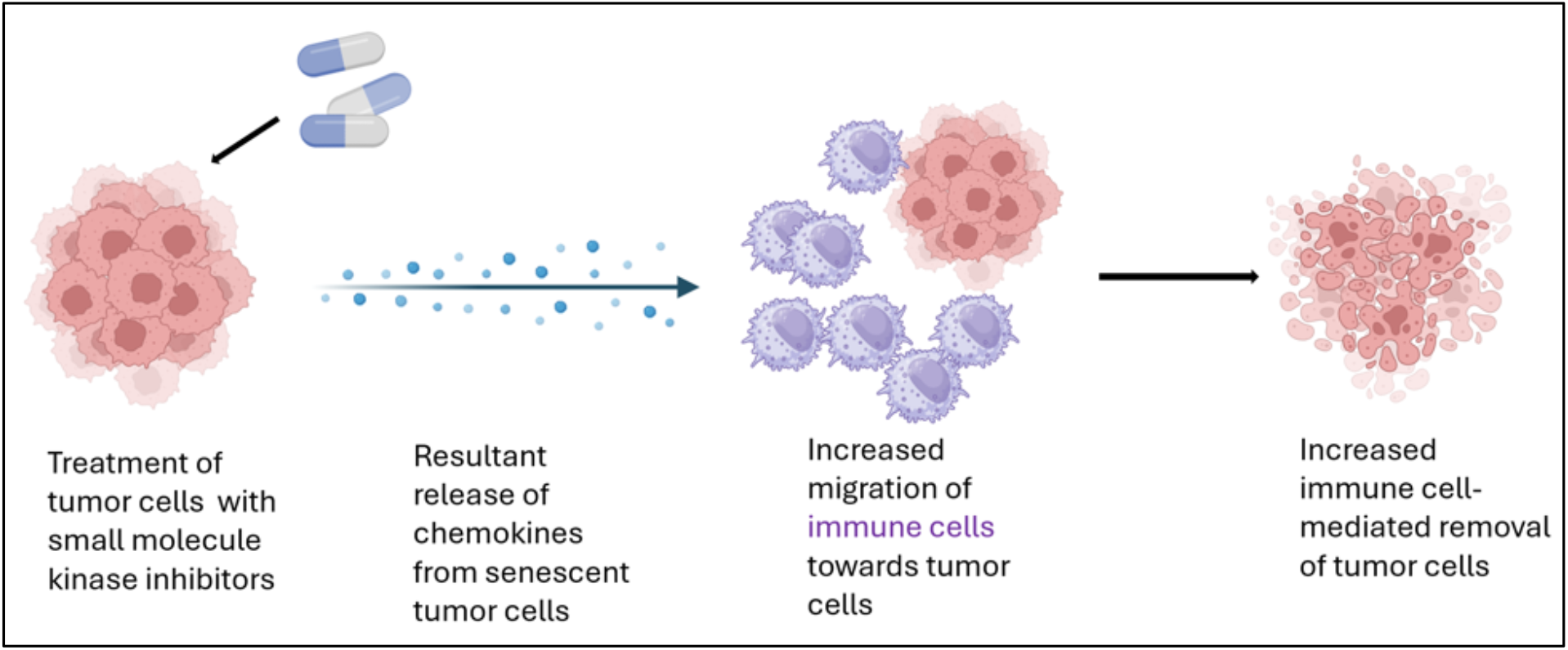

## 1. Introduction

The immune system is composed of innate and the adaptive arms, which must work in unison to protect the body foreign and/or harmful substances. Among the many different immune cell types, natural killer (NK) cells are large, granular sentinel lymphocytes. Unlike T cells, NK cells neither need priming nor depend on tumor antigen presentation. During their maturation in the bone marrow, the interaction of killer cell immunoglobulin-like receptors (KIRs) and self-composed MHC molecules ‘license’ NK cell maturation and functional capacity. Ligands for activating NKG2D receptor include MICA, MICB etc. which are often seen on cancer cells.

NK cells target cells by their MHC class I expression; a feature that most cancer cells lose during tumorigenesis. Also, unlike T cells, NK cells neither need priming nor depend on tumor antigen presentation. Thus, NK cells could potentially be deployed as an almost “off the shelf” therapeutic strategy. Moreover, target recognition of NK cells is also performed by CD16 receptor which can bind to the F_c_ region of immunoglobulins, resulting in antibody-dependent cellular cytotoxicity (ADCC), which has further contributed to the success of monoclonal antibodies like Rituximab and Transtuzumab.^1^ Now, NK cells carry out their effector functions via perforin and granzymes. Perforin can perforate the plasma membrane of target cells, creating pores and causing osmotic lysis. Granzymes (such as granzyme B) are then transferred inside to activate caspase to eventually cause target cell apoptosis.

Interestingly, the efficacy of NK cells on targeting metastatic cells in circulation or at metastatic lesions is not conclusive or mixed.^2^ Particularly, melanoma cells express a variety of ligands for various NK cell activation receptors. Additionally, NK cell-lysis of melanoma cells can depend on disease stage as well as the anatomical location due to differential expression of the ligands.^3^ Current/approved therapies for melanoma include monoclonal antibodies (mAbs) such Ipilimumab, Nivolumab, trametinib and Vemurafenib. Conventional mAbs can interact with CD16 on NK cell, thereby increasing their potential to kill cancer stem cells (CSCs) via increased antibody-dependent cellular cytotoxicity (ADCC).^4^ Radiotherapy and chemotherapy can induce cellular stress in tumor cells and thus increase NK cell-activating ligands and eventually lead to immunogenic cell death. Another abundant resident of the TME, myeloid-derived suppressor cells (MDSCs), can negatively affect NK cell function.^5^ Lastly, NK cells have found use as additional immunotherapies in combination with existing forms of chemotherapy.

These are Bi-specific and Tri-specific Killer Engagers-BiKEs and TriKEs, which are antibody reagents or small molecules with single chain variable fragments (scFv) to target the tumor antigen as well as the NK receptor. These are not antigen-specific like CAR-T cells.^6^

In melanoma (as well as in breast cancer), one therapeutic strategy is the use of small molecule kinase inhibitors like Alisertib (ALS), Abemaciclib (ABE) and Palbociclib (in breast cancer). Alisertib is known to preferentially bind to and inhibit the Aurora A kinase in cells, while Abemaciclib inhibits CDK4/6. When tumor cells are treated with these drugs, they can become senescent, entering a state in which the tumor cells do not proliferate. This strategy is used clinically in breast cancer. Interestingly, we and others observed that once in this state, the tumor cells also secrete a shifted set of cytokines and chemokines. We hypothesize that these various chemokines can attract NK cells towards the tumor cells to enhance homing to tumor cells and facilitate tumor cell killing (**as shown in the graphical abstract**). If successful, this approach could enhance NK cell homing to the proximity of tumors, allowing for NK cells to be more fully realized as an effective therapy in the clinic.

Animal models have long been the gold standard for modeling different forms of cancer. They have been important tools in many impactful scientific discoveries, but in some scenarios fail to mimic the physiological complexities associated with the disease states, as well as many nuances of human immunology. Thus, utilizing bioengineered *in vitro* models that can recapitulate these various interconnected pathways and phenomena could enable a better understanding of how to enhance NK cells function in clinical human cancer treatments. In this study, we demonstrate the use of both static 3D melanoma tumor constructs and our microfluidic device-based NK cell-functionalized tumor-on-a-chip (NK-TOC) platform as 3D microphysiological systems that can effectively be used to study homing and migration patterns of NK cells towards CDK4/6 and AURKA inhibitor treated and untreated tumor constructs in a physiologically relevant tumor microenvironment.

## 2. Results

### 2.1. Overall experimental workflow and model generation

Our approach utilized either melanoma cell lines or patient-derived melanoma cells that were encapsulated in a thiolated hyaluronic acid and methacrylated collagen-based hydrogel system, which we have utilized in past studies, which result in 3D tumor constructs (**Fig 1a-c**).^7^ Rapid crosslinking of the hydrogel biomaterials through thiol-methacrylate reactions followed by additional methacrylate-methacrylate reactions result in a hydrogel biomaterial that is user-friendly, does not require waiting for crosslinking to happen over long periods of time, and therefore supports relatively uniform distribution of cells through the 3D volume of the hydrogel. These 3D tumor constructs were successfully deployed in both Transwell-based model systems and our NK-TOC model systems to evaluate NK cell tumor homing and killing in the context of CDK4/6 and AURKA inhibition (**Fig 1d**).

**Figure 1.**
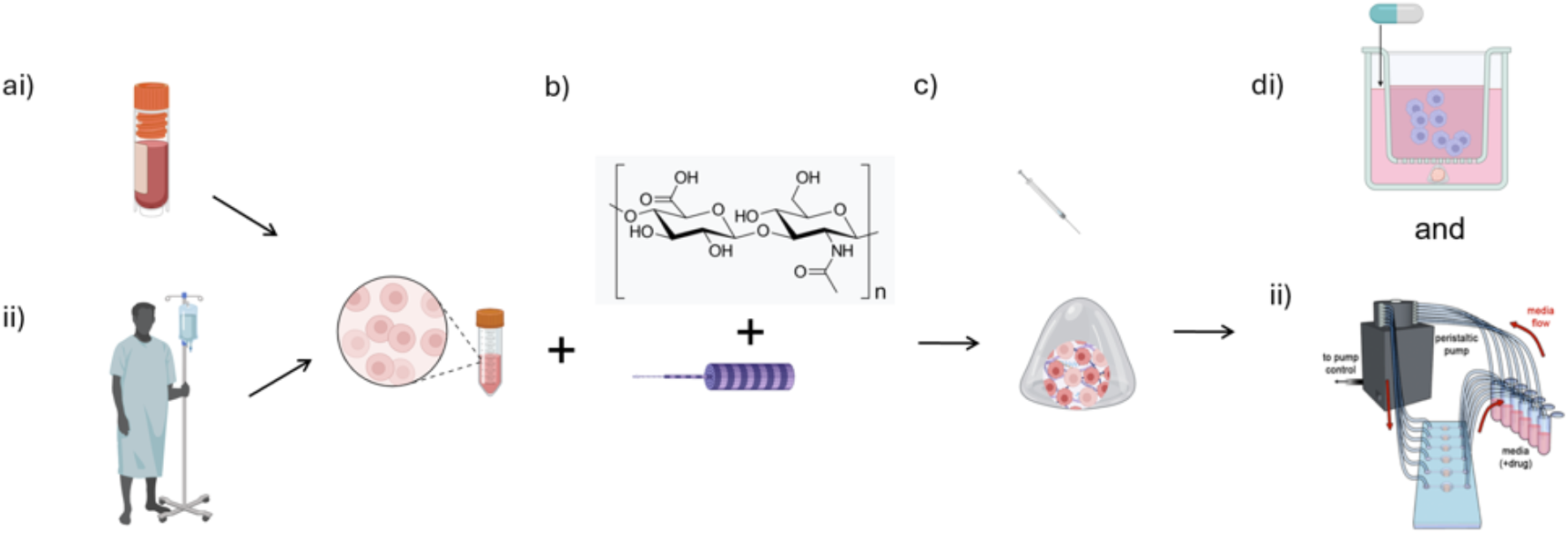
Generation of 3D models with which to study NK cell migration and effector function. (ai) Melanoma cell lines or (aii) patient-derived melanoma cells were encapsulated in (b) a hyaluronic acid and collagen-based hydrogel system to form (c) 3D tumor constructs. These 3D tumor constructs were then deployed within either (di) Transwell-based NK cell migration models and (dii) our microfluidic device-based NK cell-functionalized tumor-on-a-chip (NK-TOC) platform.

### 2.2. NK cell migration occurs within 24 hours of addition to our system and more importantly, increased NK cell migration is seen towards drug-treated samples compared to controls

Different groups have assessed migration potential of NK cells with or without additional stimulus via chemokines etc. For these migration studies, transwells or cell culture inserts are often used. The pore size of the same has varied from 0.3 μm to 5 μm.^8^ For our studies, we first used a static transwell system to test our hypothesis that kinase inhibitor treatment induces tumor cell senescence which can then express chemokines to attract NK cells and increase their migration towards tumor constructs. (**Fig 2a**). We observed increased migration of DiI-NK cells to our kinase inhibitor treated samples within 24 hours of drug treatment compared to the no drug controls (**Fig 2b**). NK cells traveled freely to all tumor organoids-drug-treated as well as control. Additionally, the cytotoxic effect of the NK cells fueled by drug treatment resulted in lower viability in these organoids compared to the control group as shown by the ATP results (**Fig 2c**).

**Figure 2.**
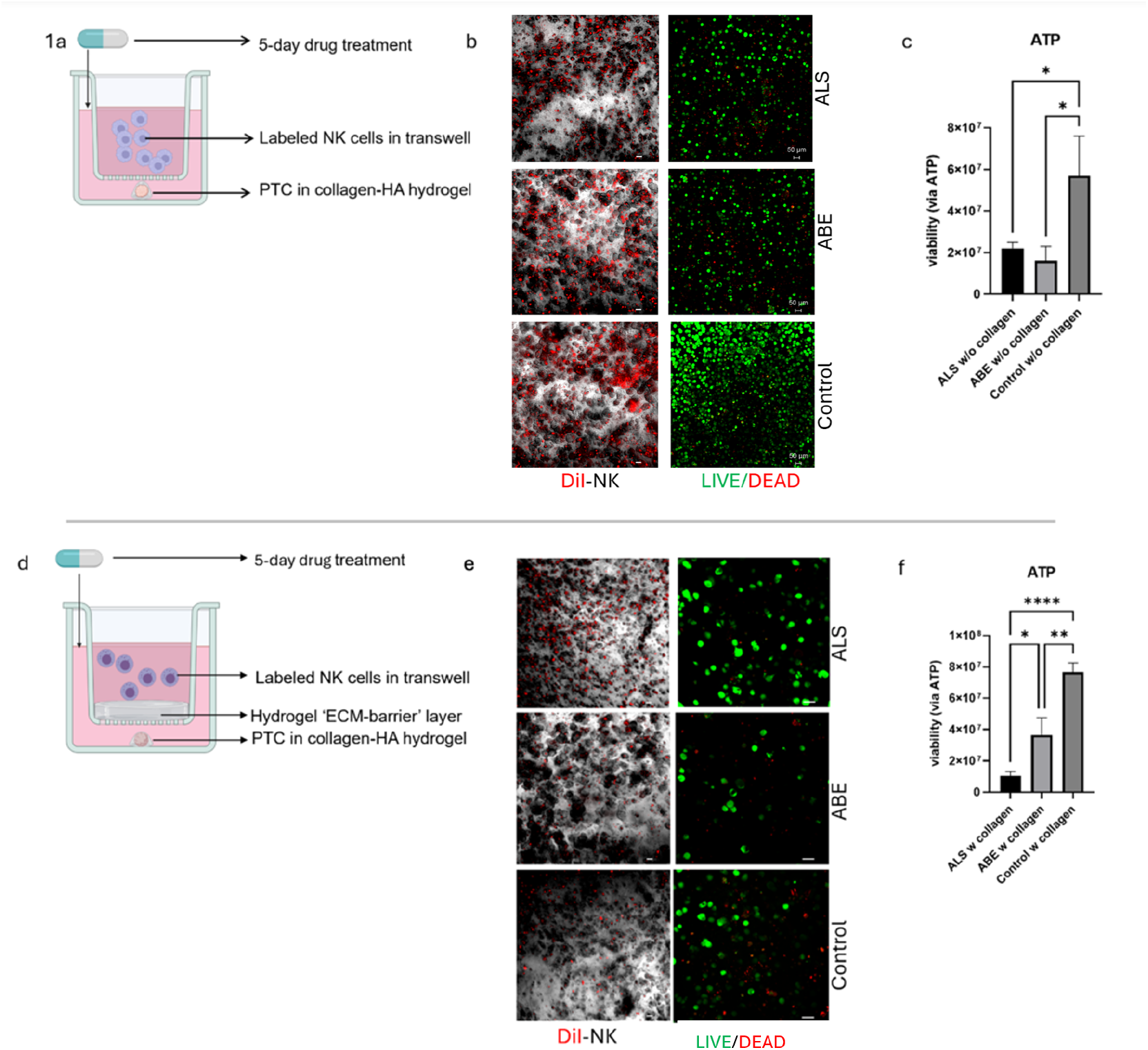
NK cell migration and viability analysis in tumor constructs in transwells. (a) Schematic of transwell phase I setup, (b) representative images for DiI-labeled NK cells confirmed migration of the same (left column) and live-dead imaging showed lower viability in drug-treated groups (right column) of tumor constructs, (c) ATP assay confirmed reduced viability in drug-treated groups compared to control. (d) Schematic of transwell phase II studies with addition of ‘ECM’ barrier layer, (e) representative images for DiI-labeled NK cells showed reduced migration of the same compared to phase I (left column) and live-dead imaging (right column) of tumor constructs showed higher viability compared to phase I results, f) viability analysis via ATP assay also confirmed lower viability across all groups compared to phase I. Green—calcein AM-stained viable cells; Red—ethidium homodimer-1-stained dead cell nuclei. Scale bar – 50 μm. Significance: * p < 0.05, ** p < 0.01, *** p < 0.001, **** p < 0.0001. Data are represented as mean + SD.

Since immune cells must travel through or cross the ECM before reaching the tumor site in vivo, we modeled this by lining the bottom of transwells with a layer of the same hydrogel to mimic an ECM ‘barrier’ (**Fig 2d**). We then measured migration of NK cells towards the tumor constructs and their corresponding cytotoxic effects. We found a clear decrease in fluorescently labeled NK cell migration through an ECM layer to underlying A-375 cell line-based melanoma organoids with Alisertib (ALS) treatment, indicating that this hydrogel ECM layer slowed NK cell movement towards the TCs. When using unlabeled NK cells for the same condition, LIVE/DEAD assays indicate increased tumor cell killing specifically in the ABE-treated condition (**Fig 2e**). Lastly, ATP results also show reduced viability in the drug-treated samples compared to the controls (**Fig 2f**).

### 2.3. Initial NK-TOC studies indicate there is higher NK cell migration to and increased effector function in kinase inhibitor treated cell line-based tumor constructs

As previously described, we first used Transwells (**Fig 2a-f**) for our studies, after which we then used a bioengineered microfluidic platform to assess NK cell homing and migration capacities. Deployment of a microfluidic platform is beneficial for several reasons. First, the glass and polydimethylsiloxane (PDMS) materials used to fabricate microfluidic devices are completely transparent, allowing for uninhibited imaging into the devices by microscopy. Second, the utilization of fluid flow adds several important features. In the context of organ-or tumor-on-a-chip systems, microfluidics allows for mimicking of the circulatory system. This enables recapitulation of shear fluid flow levels that cells might experience in the body. In our NK-TOC studies, fluid flow allows us to “challenge” the NK cells to have to exit circulation to engraft into tumor constructs.

Upon confirmation from these studies about the positive migration of NK cells induced by chemokines secreted from the kinase inhibitor-treated tumor constructs, we designed a 3D microfluidic device (**Fig 3**). Both the transwell and the device consists of have tumor constructs enclosed in ECM-mimicking hydrogels. The microfluidic device is more dynamic and challenges the NK cells to leave their normal route of circulation and migrate towards the hydrogel-enclosed tumor cells. The device has three channels-one for each condition (ALS-treated, ABE-treated and a no-drug control)-for media flow and each channel has three chambers representing tumor clusters/sites where the tumor constructs are injected. These devices can be customized to study different aspects of tumorigenesis as well as incorporate different cells and cues to activate and aid NK cell migration in a more realistic TME. Different groups have used microfluidics to evaluate not only NK cell migration in various malignancies but also other characteristics of tumor progression. ^9-15^ Our lab has also designed and used microfluidics to model various aspects of tumorigenesis. ^16-21^

**Figure 3.**
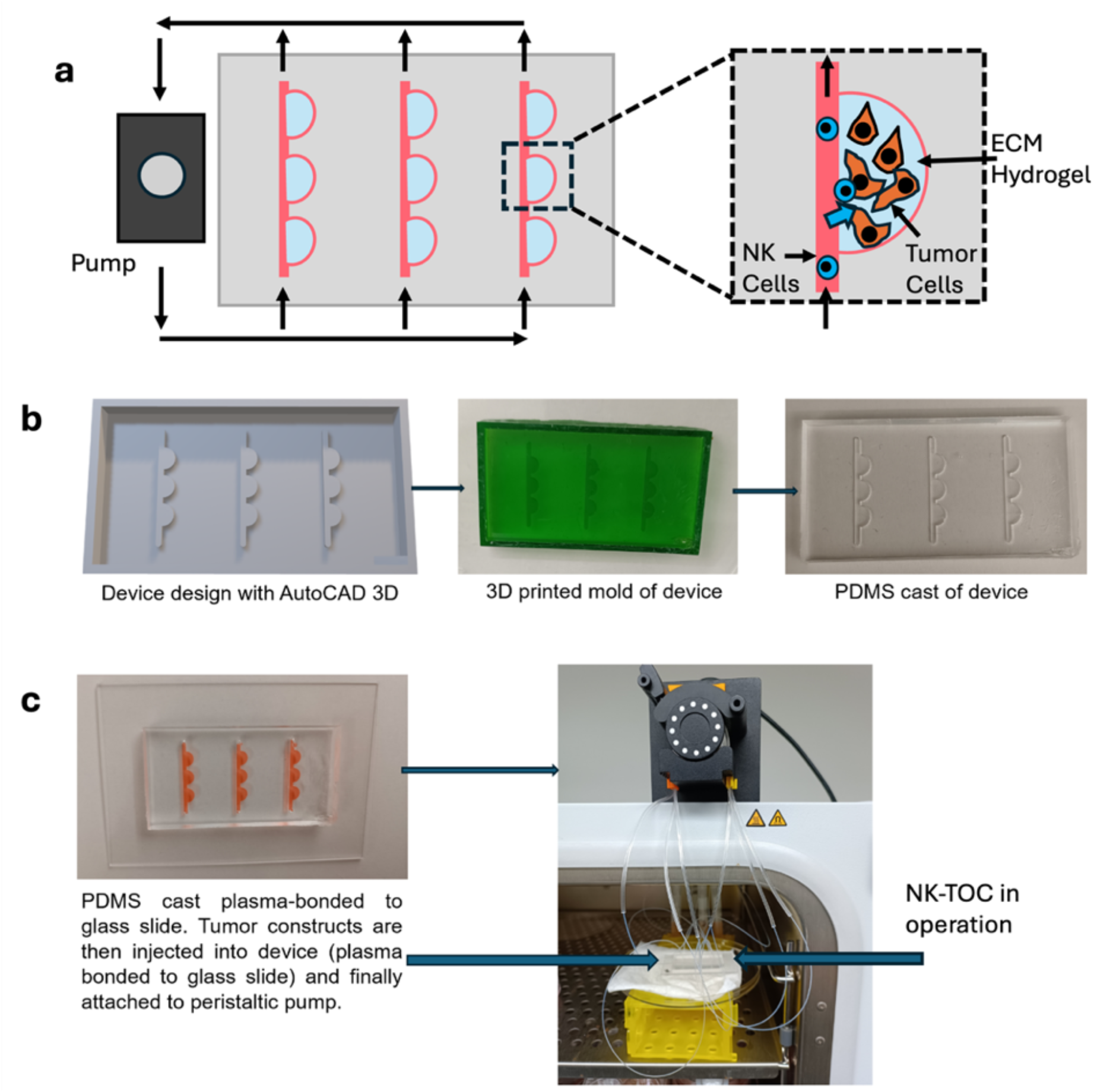
General workflow for NK-TOC device fabrication. (a) Schematic showing attachment of peristaltic pump to device and placement of tumor and NK cells in the chamber and channels respectively, (b) Device design and molding using polydimethylsiloxane (PDMS), (c) Plasma-bonded final device for attachment to pump for experiments.

We found similar patterns of NK cell migration (**Fig 4a**) in our A375 cell line-based tumor constructs from the NK-TOC platform as well. Viability/cytotoxicity measured via Live-Dead staining (and ATP assays, **Fig 4b, d**) also showed higher viability in control groups compared to organoids to which the drugs were added, especially ALS. Additionally, NK cell effector function was confirmed via Granzyme B ELISA (**Fig 4c**) and there is higher secretion of Granzyme B in ABE-treated samples compared to the no-drug control samples.

**Figure 4.**
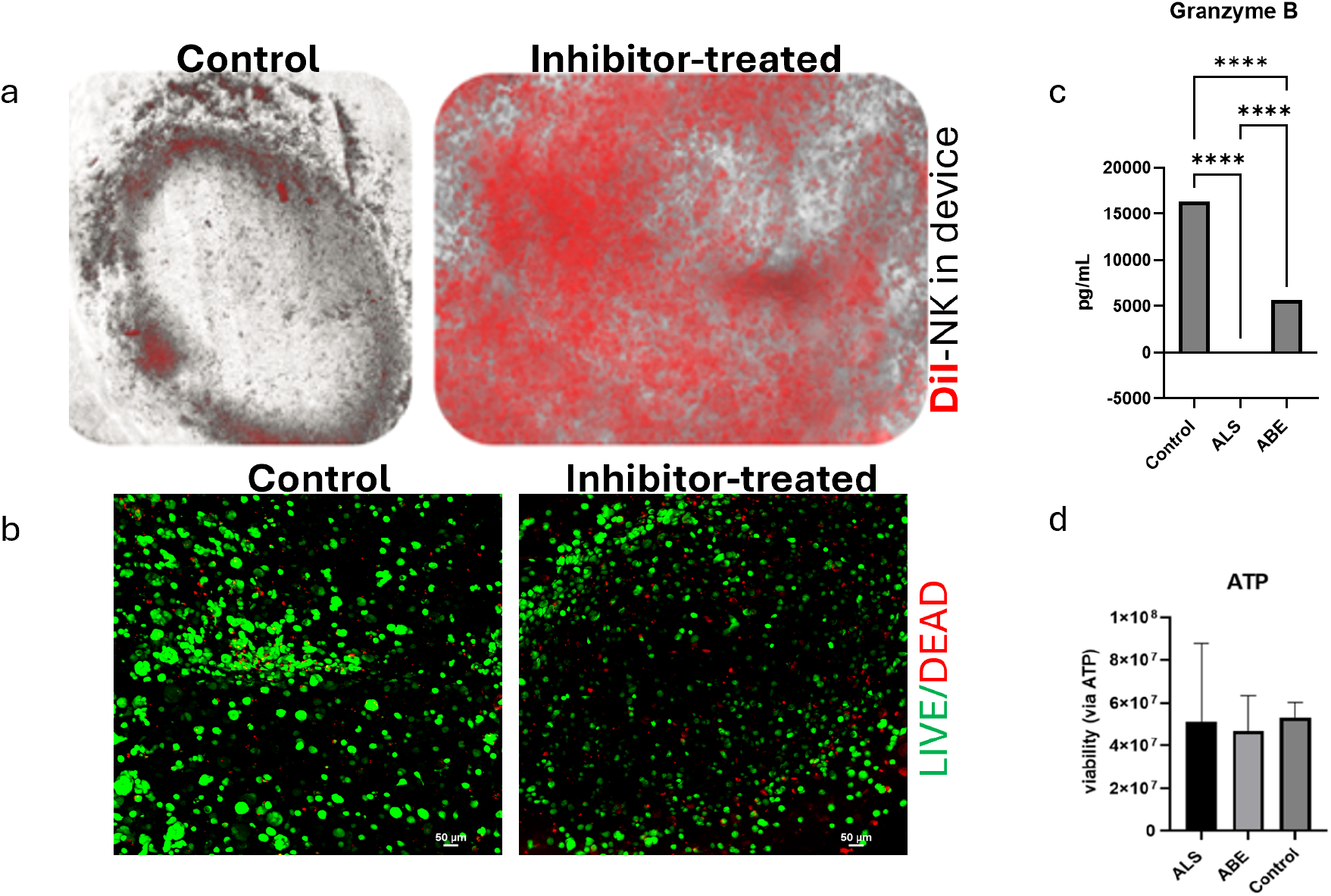
NK cell migration and viability assessment in NK-TOC platform. (a) Representative imaging of DiI-labeled NK cells showed greater migration of these cells to drug-treated, specially, ALS-treated samples (b) live-dead imaging confirmed lower viability via more red (dead) cells in ALS-treated samples (c) Expression of Granzyme B (obtained via an ELISA) validated NK cell effector function in our samples (d) viability assessment via ATP assay was not as significant. Green—calcein AM-stained viable cells; Red—ethidium homodimer-1-stained dead cell nuclei. Scale bar - 50 μm. Significance p < 0.05. * p < 0.05, ** p < 0.01, *** p < 0.001, **** p < 0.0001. Data are represented as mean + SD.

### 2.3. Kinase inhibitor treatment results in CCL5 and CXCL10 driver chemokine secretion from senescent tumor cells

ALS is an Aurora kinase A inhibitor which is known to cause senescence in different ways, including via DNA damage response which in turn is caused by increased phosphorylation of p53 among other factors. This kinase is known to control cell division via centrosome biology as well as proper spindle assembly. In different cancers, this kinase is often upregulated and the tumor suppressor function of p53 is thus downregulated. Also, p21 is a known marker of senescence and it is induced by p53 in response to the DNA damage. This in turn inhibits various cyclin-dependent kinases like CDK2 and CDK4, which can regulate cell cycle at the G1 and S phases. Additionally, p21 levels are elevated in and are associated with senescence post chemotherapy.^22^ ABE can inhibit the phosphorylation of the retinoblastoma (Rb) protein, which can cause cell cycle arrest in the G1 phase. The Rb protein is also a tumor suppressor that is phosphorylated at multiple sites during the cell cycle. This phosphorylation inactivated Rb, allowing cell cycle progression. Lastly, cancer cells often get hyperphosphorylated from the upregulation of different cyclins.^23 24-26^

Senescent marker staining for organoids from our devices showed promising results. Both CXCL10 and CCL5 were significantly elevated in drug-treated samples (particularly ALS) than in control samples (**Fig 5c-d**). CCL5 is documented to aid NK cell movement towards melanoma tumor cells and enhance autophagy-facilitated tumor cell death. CXCL10 presence is also mostly correlated with better prognosis in higher stages of melanoma. ^27-29^ For ALS-treated organoids, there was increased presence of P21 compared to controls (**Fig 5a**). Lastly, for the ABE-treated-organoids, Phospho-Rb was downregulated compared to control organoids (**Fig 5b**).

**Figure 5.**
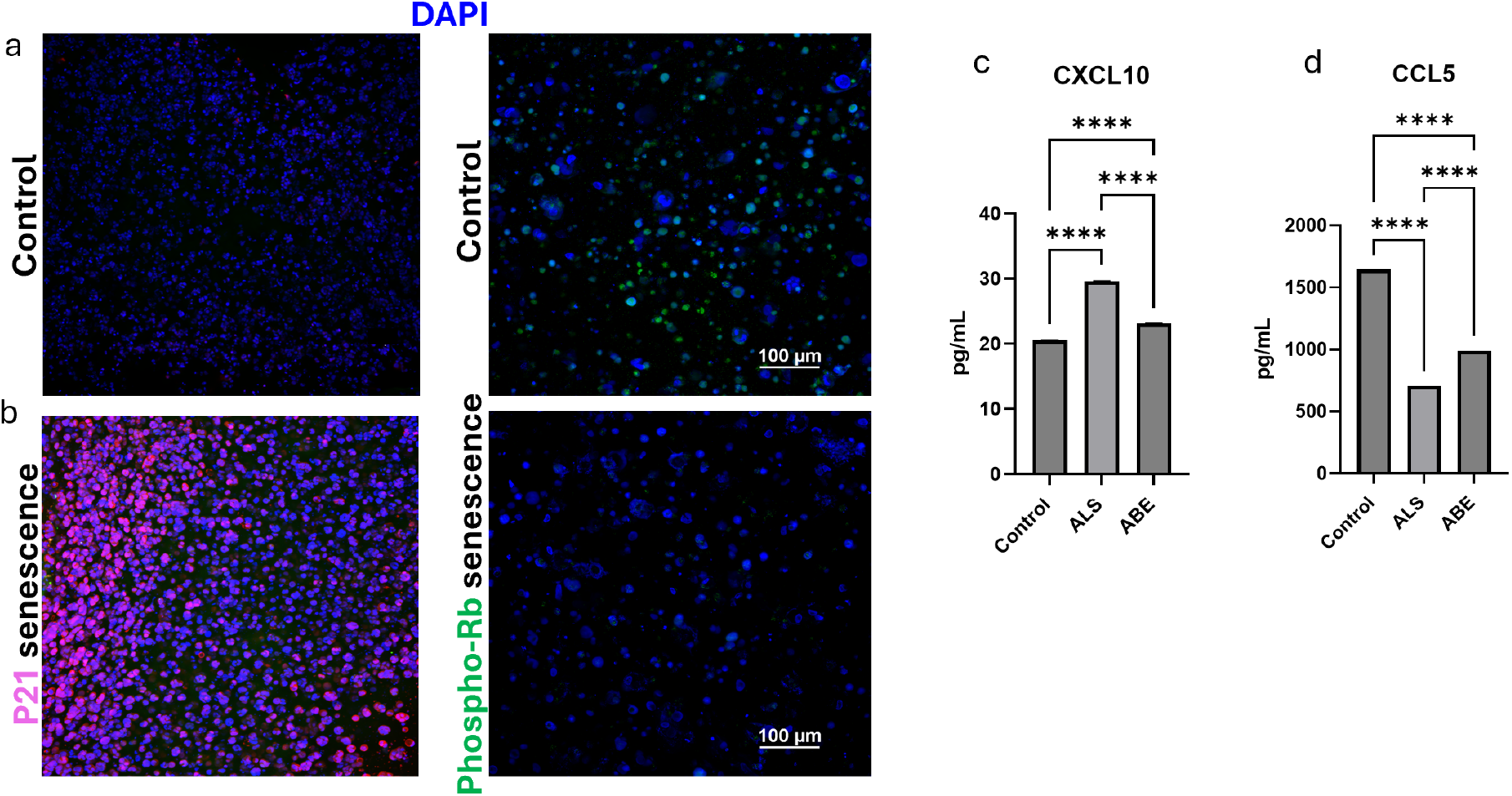
Chemokine induced senescence in NK-TOC. Representative images for P21 and Phospho-Rb senescence. (a) There is higher expression of P21 in ALS-treated samples and (b) lower expression of Phospho-Rb in ABE-treated samples compared to controls. (c-d) Driver chemokine secretion profiles via CXCL10 and CCL5 ELISAs show significant expression of these chemokines, particularly CXCL10 in ALS-treated samples compared to no-drug control samples. Scale bar - 100 μm. Significance p<0.05. *p < 0.05, **p < 0.01, ***p < 0.001, ****p < 0.0001. Data are represented as mean + SD.

### 2.6. Chemokines facilitate increased NK cell cytotoxicity in patient-derived tumor organoids

We noted higher IFN-γ and TNF-α concentrations in the drug-treated samples, further verifying action of NK cells (**Fig 6c**). NK cells are also known to use cytokines like Tumor Necrosis Factor-alpha (TNF-alpha) and Interferon-gamma (IFN-gamma)-individually or combined-to aid their cytotoxic effector function against tumor cells. Once activated, NK cells can either directly cause apoptosis of target tumor cells or employ other immune cells in the tumor microenvironment to aid in this process. However, granzyme B concentrations were inconclusive in this respect, and are hence not shown. Lastly, we also observed greater NK cell migration (**Fig 6a**) as well as greater secretion of CCL5 and CXCL10 chemokines (**Fig 6d**) in our drug-treated PDOs compared to control PDOs (PDX ID 2132 from male patient with BRAF V600E and P53 mutations). ^30^ For the drug-treated samples, specifically ALS, there is prominent organoid formation and the cells within are dead or dying off from the center to the periphery. For the control samples, there are predominantly live cells visible (**Fig 6b**).

**Figure 6.**
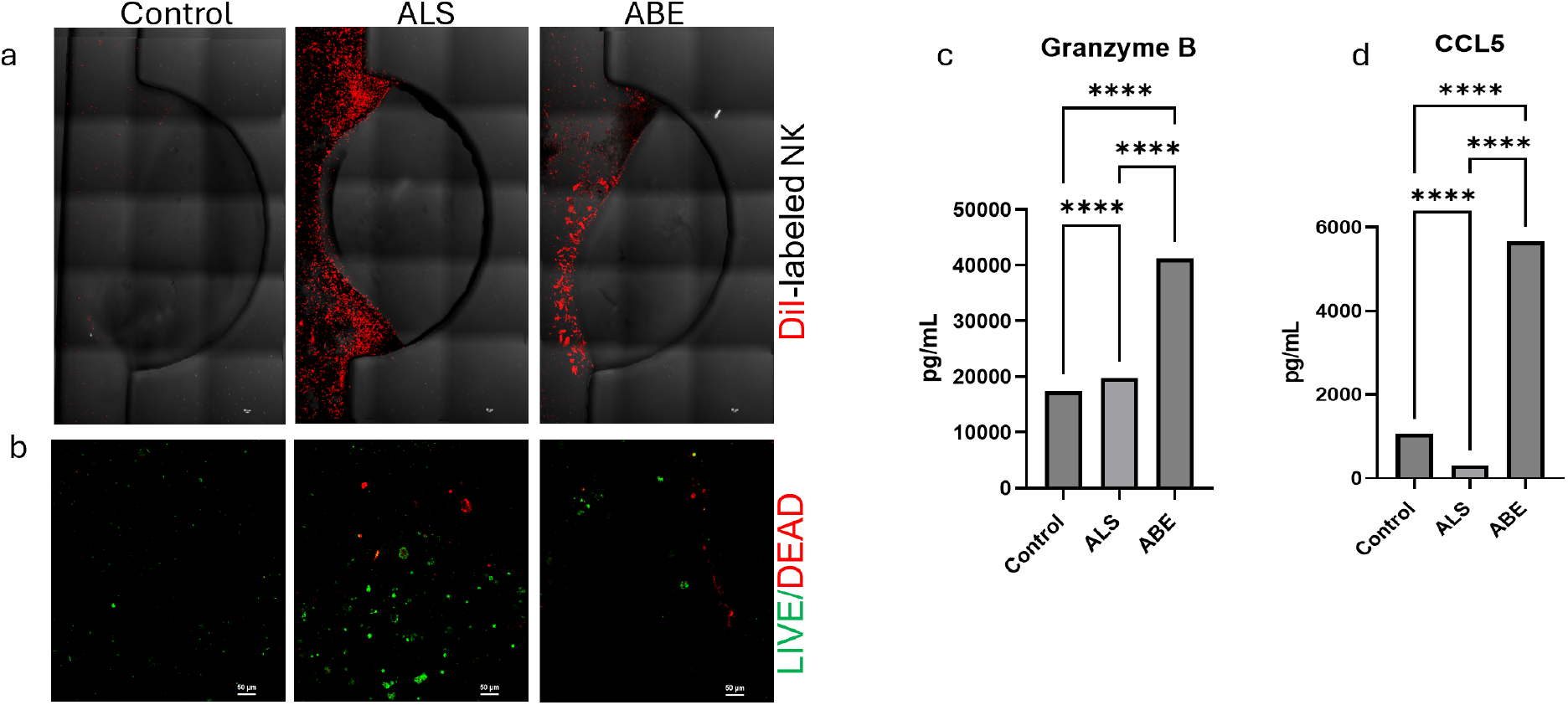
NK cell migration and viability NK-TOC platform containing patient-derived melanoma tumor organoids. (a)Representative images of DiI-labeled NK cell migration showed more effective homing of NK cells to drug-treated particularly ALS-treated groups, (b) live-dead imaging also confirms lowered viability in drug-treated samples. Reduced presence of green (live) cells in the control sample indicates higher viability, (c) NK cell effector function is validated via Granzyme B ELISA and showed significantly increased expression of these in drug-treated samples, (d) driver chemokine secretion profiles via CCL5 ELISA showed significant expression of the same in the ABE-treated samples compared to controls. Green—calcein AM-stained viable cells; Red—ethidium homodimer-1-stained dead cell nuclei. Scale bar - 50 μm. Significance p<0.05. *p < 0.05, **p < 0.01, ***p < 0.001, ****p < 0.0001. Data are represented as mean + SD.

### 2.7. Effector function of NK cells in less aggressive melanoma models is dependent on type of treatment

Similar NK cell migration and viability studies were done with mildly aggressive SK-MEL-28 cells. PTCs were constructed with these cells using the same hydrogel. Addition of labeled NK cells show a clear difference in migration patterns between the ABE-treated and control groups for the transwell setup (**Fig 7a**). For the NK-TOC platform however, there is little to no significant difference in the NK cell migration/invasion to the different groups (**Fig 7b**). Further, ELISA panels were done on media collected from these various samples to evaluate chemokine secretion as well as validate NK cell effector function. Interestingly, CCL5 levels were high only in ALS-treated samples (**Fig 7c**). Further, viability was also assessed via live-dead cytotoxicity assay (**Fig 7d**). These reflect what we observed in our migration studies in transwell models where samples with higher migration of NK cells show lower viability. But the corresponding ATP assay did not confirm this (**Fig 7e**). Now, as mentioned before, these PTCs were made of less aggressive SK-ML-28 cells and other studies using these cells document use of different kinase inhibitors and agents such as Rottlerin, CrataBL etc. This treatment inhibits proliferation of tumor cells and induces secretion of CCL5 from these cells. Also, abemaciclib has rarely been used to treat these cells and Alisertib, if used, has been used in conjunction with other drugs. ^31 32^ This could be a reason why we did not observe uniform cytotoxic effect of NK cells on the tumor cells post kinase inhibitor treatment.

**Figure 7.**
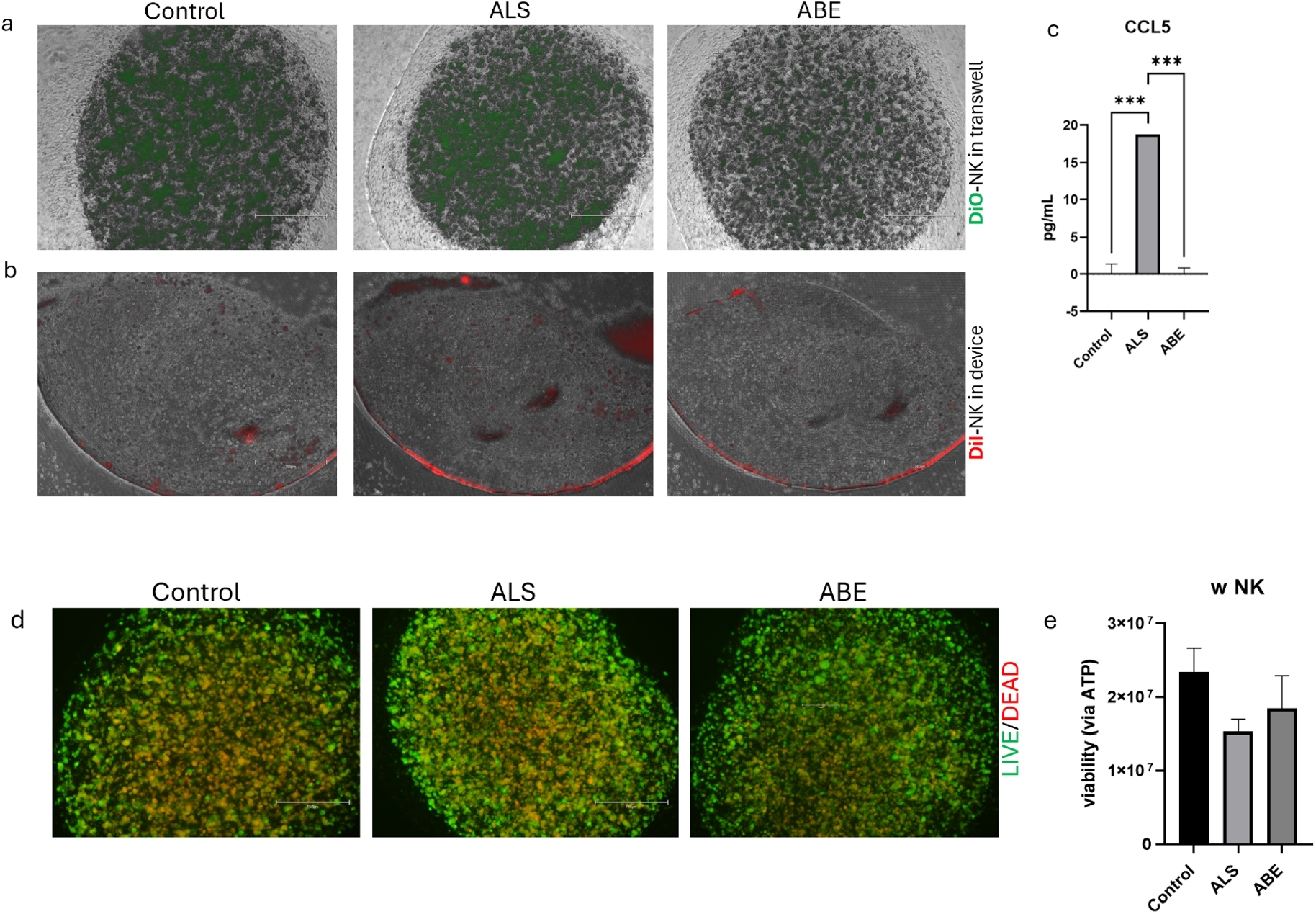
NK cell migration and viability in SK-MEL-28 tumor constructs. (a) Representative images of DiO and (b) DiI-labeled NK cell migration in transwell and microfluidic device respectively showed a visible difference in NK cell presence in the transwell but not as much in the device between the different conditions, (c) Driver chemokine secretion profiles measured via CCL5 ELISA showed differences in between the different conditions (d) viability assessment via live-dead imaging and (e) ATP assays showed reduced viability in ALS-treated samples in comparison with ABE-treated samples. Green—calcein AM-stained viable cells; Red—ethidium homodimer-1-stained dead cell nuclei. Scale bar - 750μm. Significance p<0.05. *p < 0.05, **p < 0.01, ***p < 0.001, ****p < 0.0001. Data are represented as mean +SD.

## 3. Materials and Methods

### 3.1. Cell passaging and culture

Human melanoma (highly aggressive A-375) cells and human natural killer (NK-92) cells-gifted by our collaborators at the Vilgelm lab-are expanded in 2D on tissue culture plastic in 15 cm tissue-treated dishes until 80-90% confluence. Cells are then detached from the substrate with 0.25% Trypsin/EDTA and resuspended in growth media before use in further studies. Residual A-375 cells are cultured in Dulbecco’s Modified Eagle Medium (DMEM) and 5% FBS and NK-92 cells are cultured in MEM α (Minimum Essential Medium α) till further use.

Patient-derived tumor organoids (PDOs, received from the Vilgelm lab at the Biomedical Research Tower, The Ohio State University) are first expanded from single cells to organoids in 2D 6-well plates and then used when they reach 80-90% confluency. Residual PDOs are cultured in Dulbecco’s Modified Eagle Medium/Nutrient Mixture F-12 (DMEMF12) and 15% FBS and/or frozen until needed again. Lastly, mildly aggressive SK-MEL-28 melanoma cells are expanded in 2D on tissue culture plastic in 15 cm tissue-treated dishes until 80-90% confluence. Cells are then detached from the substrate with 0.25% Trypsin/EDTA and resuspended in growth media before use in further studies. Residual cells are cultured in Eagle’s Minimum Essential Medium (EMEM) and 10% FBS till further use.

### 3.2. Cell line-based and patient-derived tumor construct preparation

ECM-mimicking collagen-hyaluronic acid (HA) hydrogels are formed using HyStem-HP (ESI-BIO, Alameda, CA). The thiolated HA (Glycosil, ESI-BIO) component is dissolved in water containing 0.1% w/v of the photoinitiator 4-(2-hydroxyethoxy) phenyl-(2-propyl) ketone to make 1% w/v solutions. Fifty-two μL of methacrylated collagen type I (Advanced BioMatrix, San Diego, CA) is added to 18.75 μL of the HA and 4.41 μL of neutralizing solution (for photocollagen) in a small conical tube. Cells are added to this resulting solution and 5 μL (Fig 10).

Tumor constructs are constructed for each well of a 48-well plate previously coated with a thin layer of polydimethylsiloxane (PDMS). For all transwell studies, tumor cell to immune cell ratio chosen is 3:1-30,000 tumor cells and 10,000 NK cells. For studies done on the devices, we use 100,000 tumor cells and 1×10^6^ NK cells.

### 3.3. Device fabrication

Microfluidic device molds are designed using AutoCAD. Molds were printed using the H-Series 3D printer (CADworks 3D, Concord, ON, CA) with CADworks 3D Mastermold Resin. The molds are used to produce devices out of polydimethylsiloxane (PDMS, Sylgard 184, DOW Chemical). The PDMS devices are left on the oven overnight to solidify and then bonded to glass slides via plasma treatment. There are inlet and outlet points in every channel for media addition. These are punched using 5mm biopsy punches. Orange-yellow two-stop tubing (0.51 mm) is attached to these channels and then to a six-channel pump (Elemental Scientific MP^2^ Stand Alone Precision Micro Peristaltic Pump) to ensure flow of media (Fig 11). These pumps allow us to control direction and velocity of flow (0.8 μL/sec and 5 rpm). The NK-TOC platform offers a more dynamic scenario and challenges the NK cells to leave their natural path of circulation to migrate and/invade into the PTCs. PTCs are injected into the device chambers using a small (0.31×8mm) insulin gauge syringe and then photopolymerized in place using a brief UV light pulse (365 nm, 18 W cm−2 for 2s to initiate crosslinking of the hydrogel.

### 3.4. Drug reconstitution and treatment

All tumor constructs are treated with 1 μM Alisertib (ALS-Selleckchem 10 mM in 1 mL DMSO) and Abemaciclib (ABE-Selleckchem 5 mg) respectively, which render the tumor cells senescent, but induce production and secretion of potent chemokines (e.g., CCL5, CXCL10 etc.) that NK cells utilize to guide migration. Both the drugs are reconstituted from their stock concentration of 10 mM to 1 μM. 0 μM is used as no-drug control/vehicle.

### 3.5. NK cell staining and flow

NK-92 cells are added on day 6. For the device, fluorescently labeled NK-92 cells (Vybrant Multicolor Cell-Labeling Kit (DiO, DiI, DiD Solutions, 1 mL each)) NK) are infused through its chambers via the pump setup. All imaging (fluorescent microscopy) and viability assays are performed on day 6 to assess NK cell homing and effector function.

### 3.6. Phenotypic and functional assays

#### 3.6.1. Viability assays

Viability was determined by LIVE/DEAD staining (LIVE/DEAD viability/cytotoxicity kit for mammalian cells; Thermo Fisher, Waltham, MA) performed 5 days post drug exposure. Spent medium is first aspirated from wells, after which 250 μL of mixture of PBS and DMEM (1:1) containing 0.5 μM calcein-AM and 2 μM ethidium homodimer-1 is introduced to each well. Tumor constructs are incubated for 30 minutes, after which fluorescent imaging is performed using a Nikon A1R confocal microscope. z-Stacks (200 μm) are obtained for each construct using filters appropriate for both red and green then superimposed. Viability is also assessed via luminescence using Cell Titer-Glo 3D cell viability assay (Promega). Spent media is aspirated from wells containing the tumor constructs and 200 μL of 1:1 mixture of 3D ATP media and DMEM is added. The well-plate is gently shaken for 5 minutes and then incubated at room temperature for 25 minutes. These contents are then transferred to a Costar White Polystyrene 96 well Assay Plate (3912, Corning, NY) and luminescence is measured on a microplate reader (Varioskan LUX 3020-80022) using default settings for luminescence protocol. The triplicate values for each condition are averaged and analyzed via using Graph Pad Prism (GraphPad, La Jolla, CA) software. Tumor constructs from the NK-TOC device are used for the same viability assays and imaged similarly.

#### 3.6.2. Protein detection assays

Media from the transwells as well as device is collected and used to detect presence of chemokines specifically CCL5 (Abcam Human RANTES ELISA Kit-ab174446) and CXCL10 (Abcam Human IP-10 ELISA kit-ab83700) via enzyme-linked immunosorbent assays (ELISA). Further, to confirm cytotoxic function of NK cells, media from these various samples is also used to detect presence of Granzyme B via ELISA. The standard protein concentrations are done in duplicates and samples in triplicates. Values are read using absorbance protocol (at 450 nm) on a microplate reader (Varioskan LUX 3020-80022). These values are then averaged, and statistical analysis is done using Graph Pad Prism software.

#### 3.6.3. Senescence detection via staining and imaging

Tumor constructs from the device are stained for senescent markers-P21 and Phospho-Rb. PTCs are removed from the device and fixed in 4% paraformaldehyde for one hour. These are then blocked and washed. Primary antibodies are thereafter added, and the PTCs are refrigerated overnight. Secondary antibodies are added after 24h and refrigerated overnight. Imaging is done on the following day. P21 (Anti-P21 antibody-Abcam ab109520) primary antibody is rabbit monoclonal and diluted at 1:1000. The secondary antibody used here is goat anti-rabbit (584 nm). Phospho-Rb used here is (Ser807/811) (D20B12) XP Rabbit mAb (Cell Signaling Technology-Alexa Fluor 488 Conjugate). All organoids-ALS-treated, ABE-treated as well as controls- are stained with both P21 and Phospho-Rb antibodies. DAPI (4′,6-diamidino-2- phenylindole, Thermo Fisher Scientific) is used as nuclear stain for all cases.

### 3.7. Statistical analysis

All samples/groups are maintained in triplicates and are averaged and analyzed via using Graph Pad Prism (GraphPad, La Jolla, CA) software. ANOVA followed by Tukey’s is used to determine statistical significance. P-value less than 0.05 is considered significant.

## 4. Discussion

In the studies described herein, we explore the use of CDK4/6 and AURKA inhibitors to prime the tumor microenvironment to be less immune-evasive to in turn allow for enhanced NK cell infiltration and tumor cell killing. While many advances in tumor immunology have been made in animal models, there are differences between murine models and humans, including in relation to metabolism and immune systems, both of which can be crucial to cancer progression and metastasis, as well as treatment efficacy. Therefore, we utilized bioengineered 3D Transwell and tumor-on-a-chip model systems within this study that were comprised of either tumor cell lines or patient-derived tumor cells. Specifically, we utilized our collagen-HA hydrogel that has been deployed across tissue engineering applications, including tumor organoids/constructs, to generate melanoma tumor constructs to test the impact of CDK4/6 and AURKA inhibitors on NK cell therapeutic intervention. Transwell-based static models were used to test feasibility after which we moved into dynamic flow-containing tumor-on-a-chip models in which infused NK cells would have to actively exit fluid flow and interact and engraft into tumor constructs.

Our results yield several interesting findings. First, our data in **Fig 1** suggests that the traditional Transwell migration assay is inadequate. The traditional Transwell migration assays do not include a mimic of ECM through which cell need to migrate, be they tumor cells, stem cells, or immune cells, regardless of the research topic. As such, without a 3D component through which cells of interest must migrate through, this assay is outdated and overly simplistic. Second, in our cell line-based Transwell and NK-TOC models we observed dramatic increases in NK cell infiltration and tumor cell killing, but mixed chemokine and cytokine changes (**Figs 2, 4**, and **5**). This suggests that we are capturing some of the complexity of tumor-immune cell interactions, but that there are likely many signals we are not capturing at this stage. Third, to address some of these limitations, we then performed NK-TOC experiments with patient-derived melanoma cells embedded within out ECM hydrogels (PTCs). These data (**Fig 6**) overwhelmingly show that CDK4/6 and AURKA inhibition significantly increase NK cell homing to the PTCs. Yet, cytokine and chemokine data is mixed, again suggesting that in future studies we will need to query additional cytokine and chemokine targets and potentially add additional cell types to our NK-TOC model to enhance cellular heterogeneity. Lastly, we performed an experiment with a melanoma cell line generally considered to be less malignant – SK-MEL-28 – as the target cell population within the NK-TOC, where results were more drug-dependent. This could potentially due to a less malignant cell line being more “immune-cold”.

Despite crucial advantages over T cells, NK cells cannot always eliminate primary tumors. Collagen type IV and laminin correspond to weakened infiltration of NK cells into primary tumors and the few NK cells that manage to reach these tumors are often poorly proliferating and weakly cytotoxic owing to lowered expression of IFN-γ. Moreover, NK cells require contact to form ‘immunological synapses’ with tumor cell to carry-out their effector functions and this can be disrupted by anti-adhesion components.^34^ Cancer cells can further gather intrinsic features such as downregulation of Fas and epigenetic changes via histone deacetylation to ultimately hamper recognition by NK cells. Also, most solid tumors such as melanoma often have hypoxic regions not only enabling metastasis but also shielding metastatic cells from immune clearance by NK cells.^35, 36^ Additionally, there is always a crosstalk between NK cells and other components of the TME such as stromal and secretory factors as well as between other immune factors.^37^ There are more than 30 types of these infiltrating stromal cells including fibroblasts which are most abundantly found.^38^ These can suppress NK cell function by downregulating the ligands of NK cell activating receptors and producing ECM components like Indoleamine-2,3-dioxygenase (IDO) and prostaglandin E2 (PGE2) as well as via other pathways.^37, 39-42^

We recognize that *in vitro* models can be quite simple, but our studies are a step towards subsequent studies that can be made more cellularly complex. Incorporation of the components from the melanoma tumor microenvironment described above can enable a realistic evaluation of the crosstalk between tumor cells, NK cells, and constituents of the TME. Our next set of ongoing studies include normal human lung fibroblasts with the melanoma cells to assess NK cell cytotoxicity and effector function.

Lastly, these studies may help us understand if we can use the same system to “train” NK cells in an immunosuppressive environment, and perhaps reduce their sensitivity to immunosuppression, generating more potent NK cells for use in other cancer types.

## Acknowledgments

A.S., A.V., and A.D. acknowledge funding from the Ohio State University Comprehensive Cancer Center (OSUCCC) and the OSUCCC Translational Therapeutics Program Seed Award. A.S. acknowledges funding from NIH grant R21 CA263137. AD acknowledges support from NIH under award number K01 OD031811. Our team acknowledges the cancer patients that consented for their excised tumor tissue to be utilized for research purposes, which led to cells and organoids that were eventually employed in the studies described herein.

